# Categorical Assignment of Pulmonary Embolism is a Simple and More Accurate Indicator of Right Ventricular Dysfunction and Short Team Mortality

**DOI:** 10.1101/642892

**Authors:** Yu Lin Chen, Colin Wright, Anthony P. Pietropaoli, Ayman Elbadawi, Joseph Delehanty, Bryan Barrus, Igor Gosev, David Trawick, Dhwani Patel, Scott J. Cameron

## Abstract

Several risk stratification tools are available to predict short-term mortality in patients with acute pulmonary embolism (PE). Right ventricular (RV) dysfunction, which is common to intermediate and high risk PE, is an independent predictor of mortality and may be a faster and simpler way to assess patient risk in acute care settings. We evaluated 571 patients presenting with acute PE as the primary diagnosis, stratifying them by the Pulmonary Embolism Severity Index (PESI), by the BOVA score, or *categorically* as low risk (no RV dysfunction by imaging), intermediate risk (RV dysfunction by imaging), or high risk PE (RV dysfunction by imaging with sustained hypotension). Using imaging data to firstly define the presence of RV dysfunction, and plasma cardiac troponin T (cTnT) and NT-proBNP as additional evidence for myocardial strain, we evaluated the PESI and BOVA scoring systems compared to categorical assignment of PE as low risk, submassive, and massive PE. Cardiac biomarkers poorly distinguished between PESI classes and BOVA stages in patients with acute PE. Cardiac TnT and NT-proBNP easily distinguished low risk from submassive PE with an area under the curve (AUC) of 0.84 (95% C.I. 0.73 – 0.95, p< 0.0001), and 0.88 (95% C.I. 0.79-0.97, p< 0.0001), respectively, and low risk from massive PE with an area under the curve (AUC) of 0.89 (95% C.I. 0.78 – 1.00, p< 0.0001), and 0.89 (95% C.I. 0.82-0.95, p< 0.0001), respectively. Predicted short-term mortality by PESI score or BOVA stage was lower than the observed mortality for submassive PE by a two-fold order of magnitude. These data suggest the presence of RV dysfunction in the context of acute PE is sufficient for the purposes of risk stratification, while more complicated risk stratification algorithms may under-estimate short-term mortality risk.

---

Pulmonary embolism (PE) is a thrombotic emergency and can be life-threatening, with an estimated incidence of > 600,000 patients annually. As the third leading cause of cardiovascular mortality following myocardial infarction and stroke, PE is responsible for approximately 200,000 deaths yearly ^1, 2^.

Various risk stratification algorithms may be utilized to determine short-term mortality following PE. The Pulmonary Embolism Severity Index (PESI) is one such algorithm used widely for predicting 30-day mortality in patients diagnosed with acute but relatively low risk PE ^3–6^. The PESI score is based on clinical variables including: sex, age, heart failure, chronic lung disease, altered mental status and malignancy as well as the patient vital signs: respiratory rate, heart rate, oxygen saturation, temperature, and systolic blood pressure < 100 mm Hg ^3, 7^ (Supplemental Figure 1). The BOVA score is also reported to predict 30-day mortality. The BOVA score is designed for use in normotensive patients but incorporates some higher risk clinical features: systolic blood pressure < 100 mm Hg, heart rate > 100 beats/min, elevated cardiac troponin (cTnT), and the presence of right ventricle (RV) dysfunction by imaging ^8^ (Supplemental Figure 2).

RV dysfunction appears to be an independent variable predicting mortality attributable to PE ^9^. The presence of RV dysfunction may increase short-term mortality by greater than two-fold in the context of acute PE ^10, 11^. RV dysfunction may be directly determined qualitatively and quantitatively by various imaging modalities, or inferred elevated using cardiac biomarkers as a surrogate. Whilst the American College of Chest physicians (ACCP) does not endorse routine assessment of plasma cardiac biomarkers in every patient diagnosed with PE, the European Society of Cardiology (ESC) guidelines include the PESI score as well as cardiac biomarkers as well as cardiac imaging information to further risk stratify a patient with acute PE ^12–14^. Recent studies suggest that elevated plasma cardiac troponin in patients with PE is associated with an increase in all-cause mortality^12^.

The use of Pulmonary Embolism Response Teams (PERTs) to rapidly assess patient risk and to institute a treatment algorithm for patients with submassive and massive PE has gained popularity around the world, and may improve both mortality and the efficiency of treatment in acute care environments such as the emergency department ^15–18^. Prognostic information obtained during expedited patient evaluation may influence the choice of advanced treatment options and recently was reported as an independent variable predicting mortality ^19^. The PESI score and BOVA stage are used by many clinicians to predict short-term mortality, but may be misleading in the context of a PERT evaluation where patient acuity is higher. The BOVA score documents the presence of RV dysfunction directly (imaging) and indirectly (elevated cardiac troponin) while the PESI score relies on multiple other clinical variables.

A much simpler way to categorize acute PE for the purposes of a PERT evaluation may be the presence of RV dysfunction: 1. Low risk PE (normotensive, no RV dysfunction), 2. Intermediate risk or “submassive” PE (normotensive with RV dysfunction), 3. High risk or “massive” PE (SBP < 90 mmHg for at least 15 minutes attributable to RV dysfunction) ^20^. To illustrate this point, Sanchez *et al*. reported that RV dysfunction by either echocardiography or CTA, or by elevated plasma cardiac biomarkers predict increased mortality — even in patients with hemodynamically stable PE ^21^.

The present investigation examined individuals presenting to a single center with a diagnosis of acute PE. We used imaging and blood pressure to stratify PE into low risk, submassive, or massive categories. We then assessed cardiac biomarkers as indirect surrogates of RV dysfunction. The goal was to ascertain whether cardiac biomarkers, often obtained immediately on patient arrival, are useful in distinguishing between PESI classes, BOVA stages, and PE stratified categorically as low risk, submassive, and massive. We also assessed whether short-term mortality predicted by the PESI score and BOVA stages was accurate and provided additional meaningful information.

## Methods

This is a single center retrospective analysis of adult patients during a twenty-month period (May 2014 to December 2015) with a diagnosis of “pulmonary embolism” or “venous thromboembolism” based on International Classification of Diseases, 9^th^ Clinical Modification (ICD-9-CM) codes. The study protocol, data collection and storage were approved by University of Rochester Institutional Review Board. Patients were categorized into low risk PE, submassive PE, or massive PE groups and their respective PESI classes and BOVA based on score. PESI scores were calculated and grouped in three stages: PESI I (< 65 points), PESI II/III (66-105 points) and PESI IV/V (>106 points) (Supplemental Figure 1) ^22^. BOVA scores were documented as BOVA stage I (0-2 points), BOVA stage II (3-4 points, BOVA stage III (> 4 points) (Supplemental Figure 2) ^8^. We defined the diagnosis of <u>low risk</u> PE as normotensive and the absence of documented RV dysfunction by imaging. <u>Submassive PE</u> was defined as RV dysfunction by computed tomography angiography (CTA) or echocardiography (Supplemental Figure 3). <u>Massive PE</u> was defined as hypotension with a systolic blood pressure < 90 mm Hg for at least 15 minutes or cardiopulmonary arrest or cardiogenic shock by clinical exam.

Data collected included the initial set of plasma cardiac biomarkers (NT-proBNP, cardiac troponin), objective documentation of RV function by imaging, and mortality at 1 month, 3 months, and 6 months. Cardiac biomarker data was used to distinguish between low risk PE, submassive PE, and massive PE as well as between PESI classes and BOVA stages using indicated by Receiver Operator Characteristic (ROC) curve analysis. ROC curve data were reported as the area under the curve (AUC), along with sensitivity and specificity and biomarker concentration cut-points. It is important to note that the ROC curves used for categorically assigned classes of PE were based on RV dysfunction by imaging which allowed us to fairly assess the predictive ability of cardiac biomarkers in acute PE. Dichotomous variables are presented as frequencies and continuous variables as mean with standard error of mean (SEM) unless otherwise stated. The distribution of data was interrogated for normality using the Shapiro–Wilk test before comparison between groups. For data that were not Gaussian-distributed, the Mann-Whitney *U* test was used when comparing two groups, and the Kruskal–Wallis test followed by Dunn post-test was used for three or more group comparisons. For Gaussian-distributed data, the student’s t-*test* was used to compare two groups and, for three or more group comparisons, 1-way ANOVA then the Bonferroni multiple comparisons test was used. Significance was determined if the P value was < 0.05. All data were analyzed with GraphPad Prism 7 (GraphPad Software, Inc, La Jolla, CA).

## Results

### Study Population

571 patient charts were reviewed, 286 of which were determined to be clinical presentations of acute PE. Patients were excluded following chart review if a primary diagnosis other than PE was determined, if the diagnosis was made at another institution, or if the patient had only a history of VTE/PE (Figure. 1). The demographic variables and observed short-term outcomes for the patient population are shown in Table 1.

**Fig. 1:**
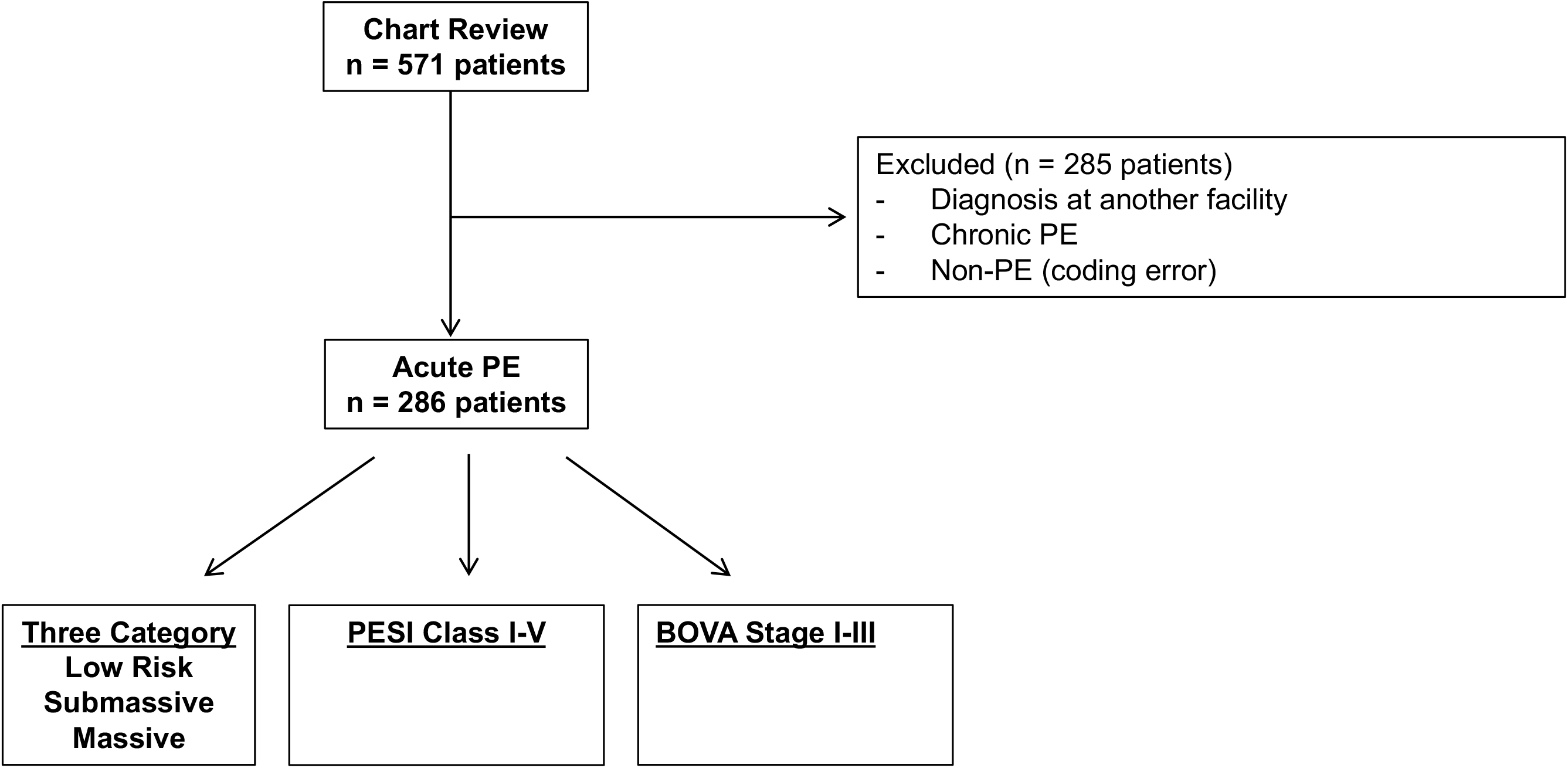
Inclusion criteria for patients screened by ICD-9 Code.

**Table 1:** Baseline Characteristics of Patient Population.

| PE Classification | Low risk | Submassive | Massive |
| --- | --- | --- | --- |
| N (%) | 149 (52.1) | 108 (37.8) | 29 (10.1) |
| Sex, n (M/F) | 83/66 | 60/48 | 14/15 |
| Race (C/O**)(%) | 105/44 (70.5) | 76/32 (70.4) | 14/15 (48.3) |
| Age (years ± SEM) | 54.0 ± 1.4 | 62.9 ± 2.9 | 62.0 ± 2.6 |
| Smoker (Y/N) (%) | 79/70 (53.0) | 59/49 (54.6) | 14/15 (48.3) |
| Provoked PE (%) | 105/149 (70.5) | 72/108 (66.7) | 21/29 (72.4) |
| Cancer (%) | 48/105 (45.7) | 31/72 (43.1) | 8/21 (38.1) |
| Chronic Lung Disease (%) | 19/149 (12.7%) | 28/108(25.9%) | 3/29 (10.3) |
| **Mortality , n (%)** |  |  |  |
| 30d | 6 (4.0) | 9 (8.3) | 9 (31) |
| 3 month | 16 (10.7) | 21 (19.4) | 11 (37.9) |
| 6 month | 23 (15.4) | 25 (23.1) | 12 (41.4) |

### Plasma Cardiac Biomarker Concentration in patients with acute PE according to PESI Class or Categorical Assignment

We used imaging data to determine the presence of RV dysfunction. We then examined categorical assignment of PE severity, observing a stepwise increase in plasma cTnT and NT-proBNP from low risk PE to massive PE. Conversely, using the PESI scoring system, an upward trend in plasma cardiac biomarker concentration was not clearly observed with ascending PESI class (Figure 2).

**Fig. 2:**
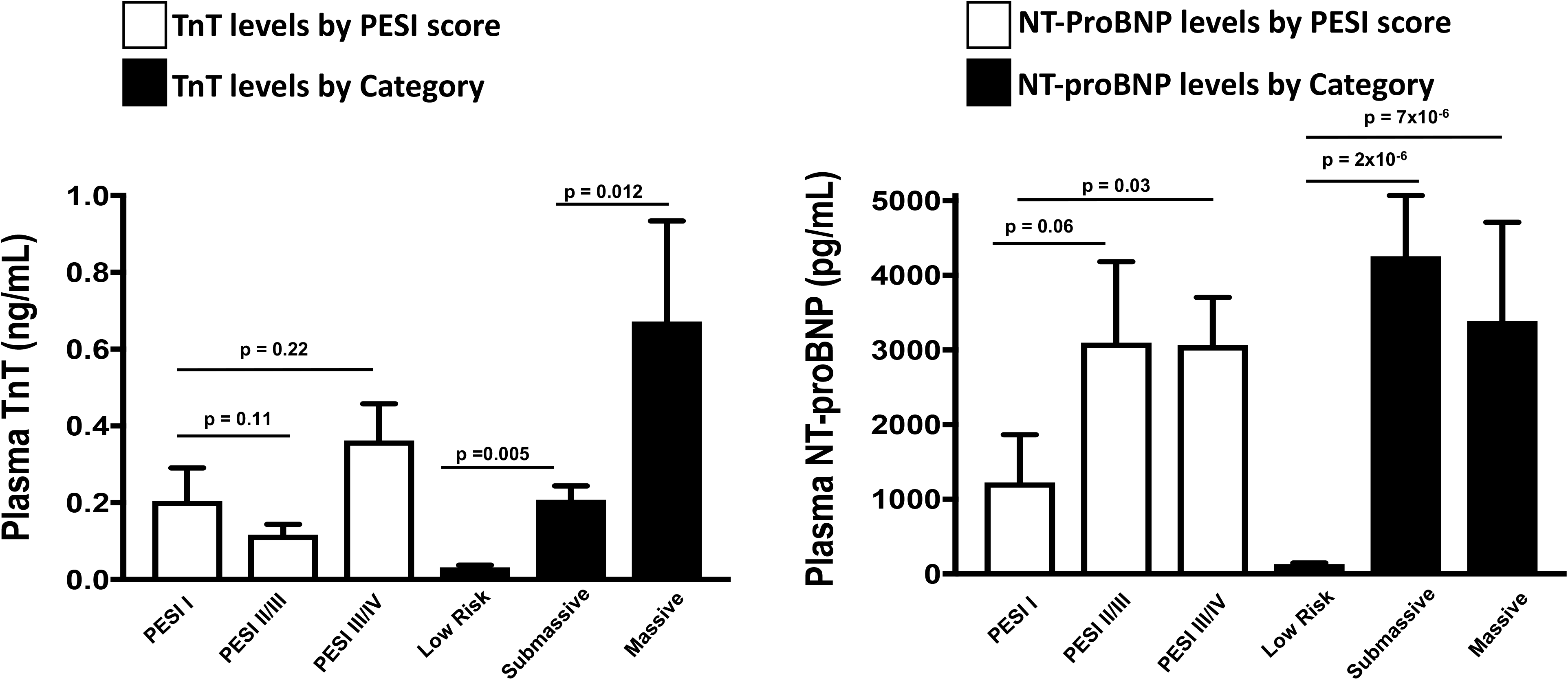
Plasma cardiac biomarker concentration of PE by categorical assignment as low risk PE, intermediate risk (submassive) PE, or high risk (massive) PE. Patients were also stratified according to the Pulmonary Embolism Severity Index (PESI) score I through V. Where available, the first plasma biomarker concentration was reported for plasma cardiac troponin and NT-proBNP as mean ± SEM. TnT=cardiac troponin T. NT-proBNP=N-terminal pro-brain natriuretic peptide.

### Performance of Cardiac Biomarkers in patients with acute PE for distinguishing between PESI Class compared to Categorical Assignment

Based on the ROC analysis of cardiac biomarkers in patients with RV dysfunction determined purely by imaging, cTnT distinguished between low risk PE and submassive PE with AUC 0.84 (95% C.I. 0.73 – 0.95). Similarly, TnT distinguished between low risk PE and massive PE with AUC 0.89 (95% C.I. 0.78 – 1.00). For NT-proBNP, low risk PE was distinguished from submassive PE with AUC 0.88 (95% C.I. 0.79 – 0.97), and low risk PE was distinguished from massive PE with AUC 0.89 (95% C.I. 0.78 – 1.0) (Figure 3).

**Fig. 3:**
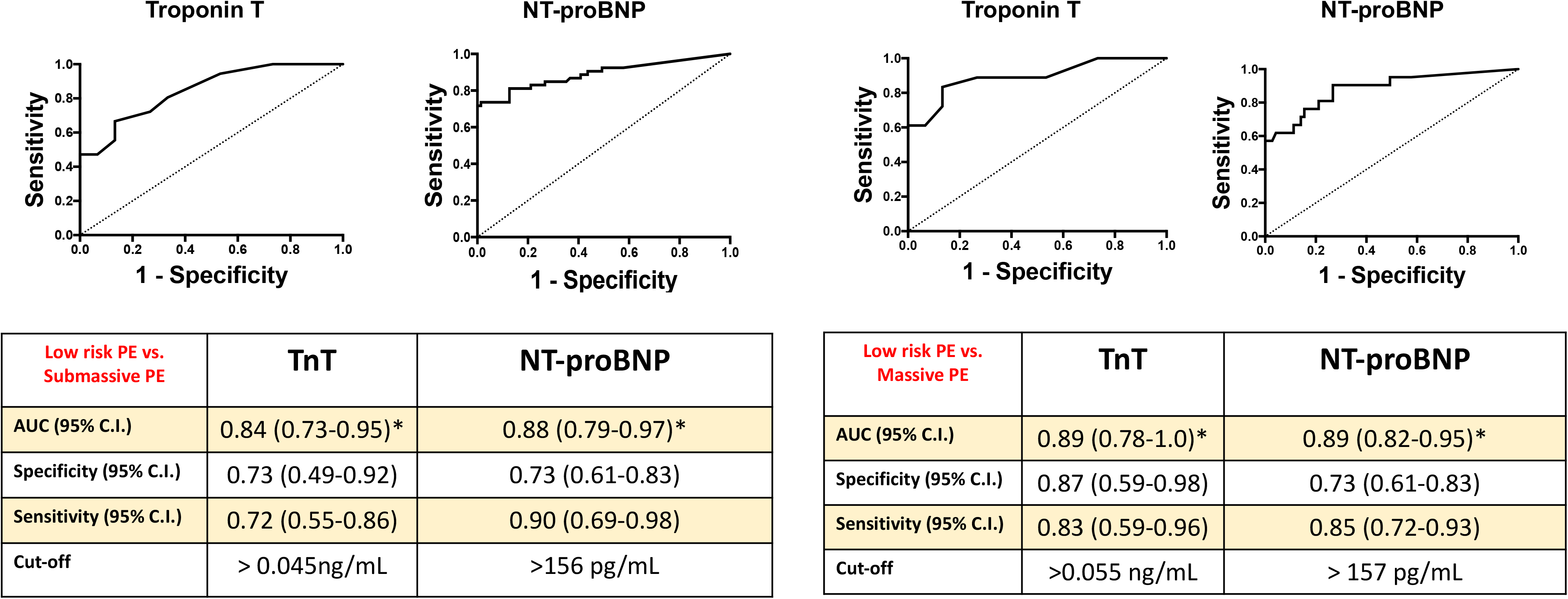
Plasma cardiac biomarker prediction of PE by categorical assignment as low risk PE, intermediate risk (submassive) PE by RV dysfunction on imaging, or high risk (massive) PE. The performance of each biomarker in predicting submassive and massive PE was evaluated by Receiver Operator Characteristic (ROC) Curve analysis for specificity, sensitivity, area under curve (AUC), and cut-off shown. The dashed line is the line of identity. C.I.=confidence interval. TnT=cardiac troponin T. NT-proBNP=N-terminal pro-brain natriuretic peptide.

In striking contrast, cTnT distinguished poorly between all PESI classes: PESI I vs. PESI II/III with an AUC of only 0.52 (95% C.I. 0.28 – 0.73), and PESI I vs. IV/V with an AUC of only 0.55 (95% C.I. 0.31 – 0.76). NT-proBNP was similarly less precise in distinguishing among the severity of PESI classes, with an AUC of 0.68 (95% C.I. 0.57 – 0.79) for PESI I vs. II/III, and an AUC of AUC 0.77 (95% C.I. 0.69 – 0.85) for PESI I vs. IV/V (Figure 4).

**Fig. 4:**
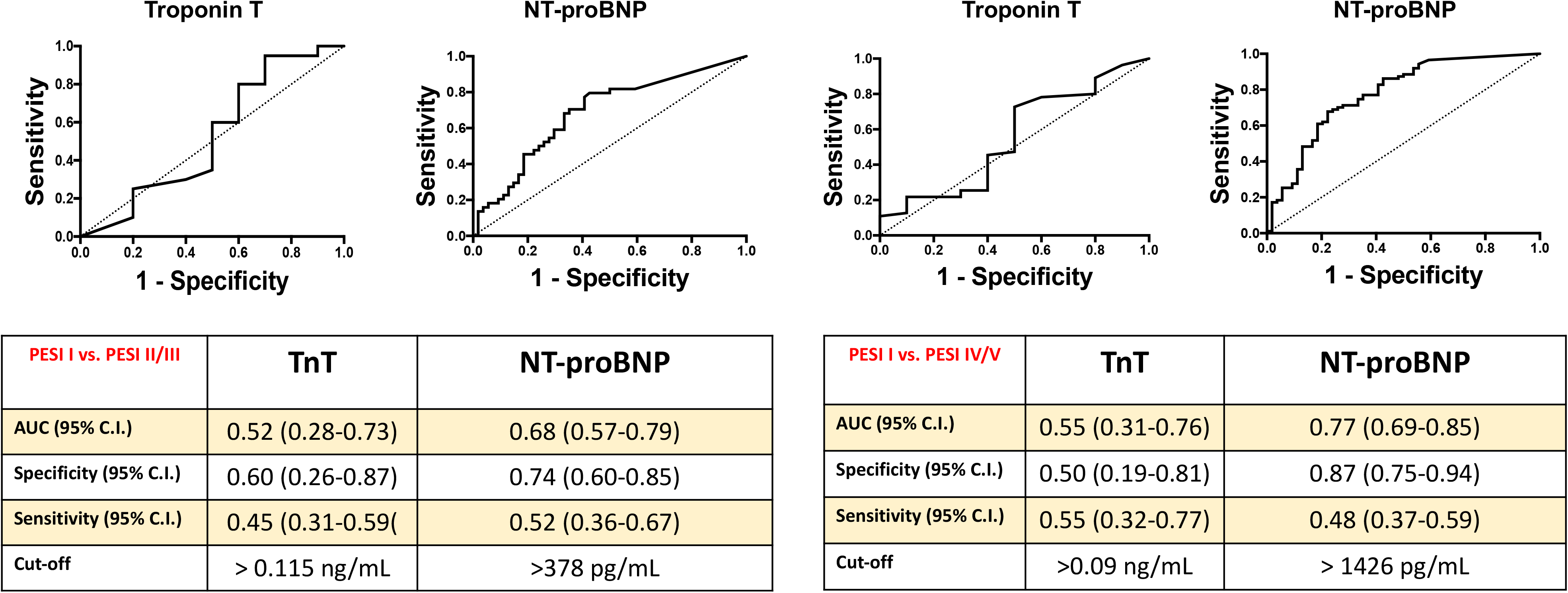
Plasma cardiac biomarker prediction of PE by PESI Scores class I through V. The performance of each biomarker in predicting low risk, intermediate risk (submassive) and high risk (massive) PE was evaluated by Receiver Operator Characteristic (ROC) Curve analysis for specificity, sensitivity, area under curve (AUC), with cut-off shown. The dashed line is the line of identity. C.I.=confidence interval. *p< 0.0001 between groups. PESI=Pulmonary Embolism Severity Index.

Using the categorical classification of low risk, submassive, or massive PE, the mean calculated PESI scores trended in an ascending manner. However, when attempting to determine a PESI score cut-point by ROC curve analysis, the value obtained to distinguish low risk from submassive PE was 93 (30 day predicted mortality of 4.8%, Figure. 5) which was 1.8-fold lower than the observed mortality (Table 1). Similarly, the predicted PESI score cut point to distinguish low risk PE from massive PE by ROC curve analysis was 108 (30 day predicted mortality of 4.8%) which was 7.2-fold lower than the observed mortality for massive PE (Table 1).

**Fig. 5:**
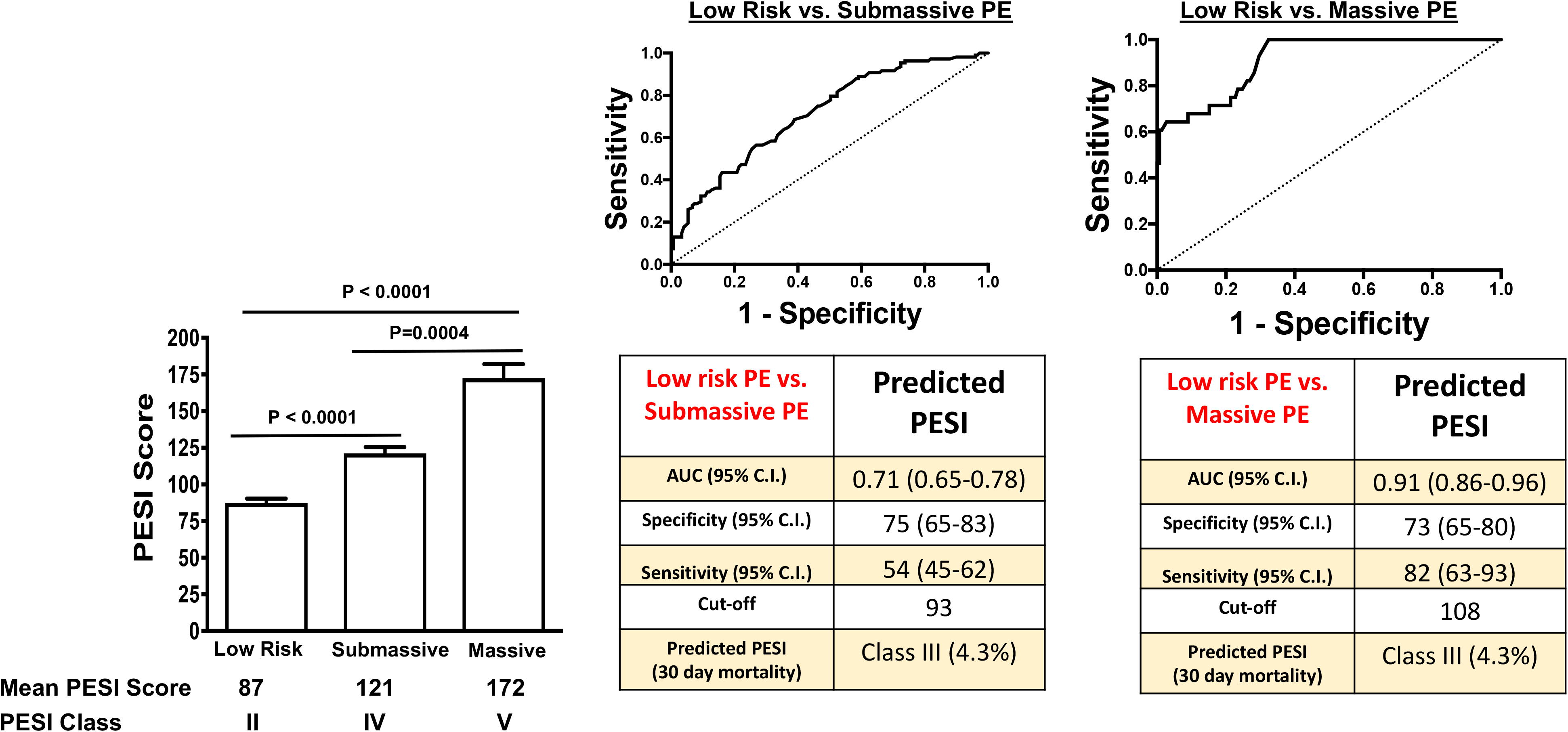
A PESI Score cut-point was calculated to distinguish between patients with low risk PE and intermediate risk (submassive) PE or high risk (massive) PE by Receiver Operator Characteristic (ROC) Curve analysis for specificity, sensitivity, area under curve (AUC), with cut-off shown. The dashed line is the line of identity. C.I.=confidence interval. *p< 0.05 between groups. PESI=Pulmonary Embolism Severity Index.

**Fig. 6:**
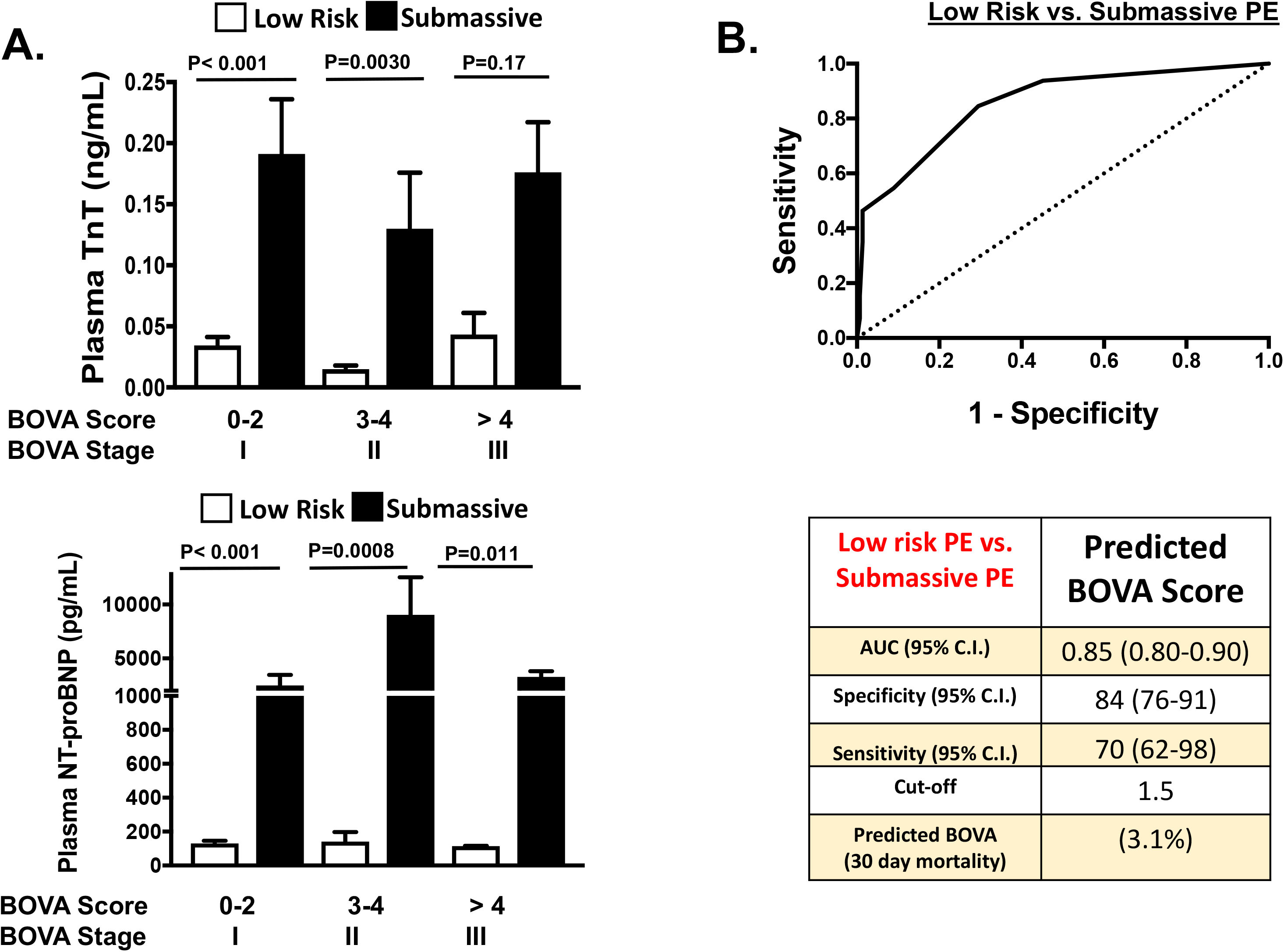
*A*, Plasma cardiac troponin concentration was assessed in patients determined to have low risk PE or intermediate risk (submassive) PE according to BOVA stages I through III. *B*, A BOVA Score cut-point was calculated to distinguish between patients with low risk PE and intermediate risk (submassive) PE by Receiver Operator Characteristic (ROC) Curve analysis for specificity, sensitivity, area under curve (AUC), with cut-off shown. The dashed line is the line of identity. C.I.=confidence interval.

### Performance of Cardiac Biomarkers in patients with acute PE for distinguishing between BOVA stages compared to Categorical Assignment

Whilst the BOVA score was validated as a short-term predictor of mortality in patients with normotensive (low risk and submassive) acute PE ^8^, components of BOVA (Supplemental Figure 2) do include certain high-risk features such as RV dysfunction and elevation of a plasma biomarker of myocardial necrosis. In spite of these features of BOVA, all BOVA stages showed markedly lower plasma cTnT concentration compared to patients who were stratified into the submassive PE category in each BOVA stage based only on radiographic evidence of RV dysfunction. Furthermore, assessing the BOVA score for each patient with low risk PE compared to submassive PE, ROC curve analysis revealed a BOVA stage cut-point predicting a 30-day mortality of 3.1%, which was 2.7-fold lower than the observed mortality in our patients with submassive PE (Table 1)

Together, these data imply that a biomarker of myocardial strain, a biomarker of myocardial necrosis, and the presence of RV dysfunction radiographically—with or without sustained hypotension—in submassive and massive PE, respectively, provide more accurate information for patient mortality risk not accounted for by the PESI or the BOVA score.

## Discussion

According to published registries, the average 3-month mortality for a patient with submassive PE is 20-25% and 50-60% for a patient with massive PE ^20^. Therefore, the presence of RV dysfunction or sustained hypotension are simple and more relevant clinical variables to rapidly determine patient risk irrespective of whether a hospital utilizes the PERT concept to assist in immediate risk stratification. As proof of concept, we calculated the PESI score and BOVA score for patients shown to have low risk, submassive, or massive PE. Our investigation confirmed that both PESI and BOVA underestimate short-term mortality for patients with acute PE in the setting of RV dysfunction or sustained hypotension. We further substantiated this observation by showing cardiac biomarkers suggestive of necrosis (cTnT) and strain (NT-proBNP) track more reliably with categorical assignment of a patient as low risk, submassive, or massive PE.

Cardiac troponin can be released from necrotic myocytes in the setting of ischemia as well as both acute and chronic myocardial loading conditions. BNP and its inactive N-terminal fragment NT-proBNP, unlike atrial natriuretic peptide (ANP) or cTnT, are not present in healthy cardiac myocytes in large quantities, and require chronic myocyte stretch to be synthesized and stored as myocyte granules ready for secretion ^12, 23–28^. We postulate the time needed to synthesize and release granules containing BNP accounts for the more impressive increase in plasma NT-proBNP concentration observed in patients with submassive PE—many of whom likely experience a subacute or chronic increase in myocardial load prior to evaluation. When cardiac biomarkers were evaluated by PESI classes or BOVA stages, we found no reliable association between plasma cTnT concentration and calculated scores. Published data indicate not only plasma troponin but also natriuretic peptides are predictive of adverse patient outcomes in the setting of PE, which is a variable not accounted for by the PESI or BOVA stage ^29–31^.

Our investigation has several limitations. Consistent with previous studies of patients with acute PE in whom RV dysfunction was documented, there is a subjective component to this interpretation, particularly when echocardiography is the imaging modality utilized. In addition, while a measured RV/LV ratio > 0.9 in the apical 4 chamber view on either CTA or echocardiography suggests RV dilation, volume status, and chronic medical conditions such as obesity, pulmonary hypertension, and right-sided valvular insufficiency all augment RV loading conditions. Therefore, any cause of increased contrast noted in the pulmonary vasculature and RV by CTA is subject to being interpreted as evidence on “RV dysfunction”, as reported ^20, 32–34^. Our study also excluded patients with a pre-existing diagnosis of VTE/PE, or patients with diagnosis of acute PE at an outside facility who were later transferred in to our institution. These factors limit external validity.

## Conclusions

Using cardiac biomarkers as a surrogate for RV dysfunction, which was determined separately by imaging or sustained hypotension, we found that the PESI and BOVA scoring systems underestimate a patient’s true short-term mortality risk in the clinical context of acute PE. We therefore suggest that when a rapid decision is required to assess patient risk or to select a treatment plan in the context of acute PE—which is the nature of a PERT—the presence of RV dysfunction and elevated cardiac biomarkers are the most relevant clinical variables determining short term mortality.

## Acknowledgements

We thank Peter Wilmot for assisting in study implementation.

---

**Supplemental Figure 1:**
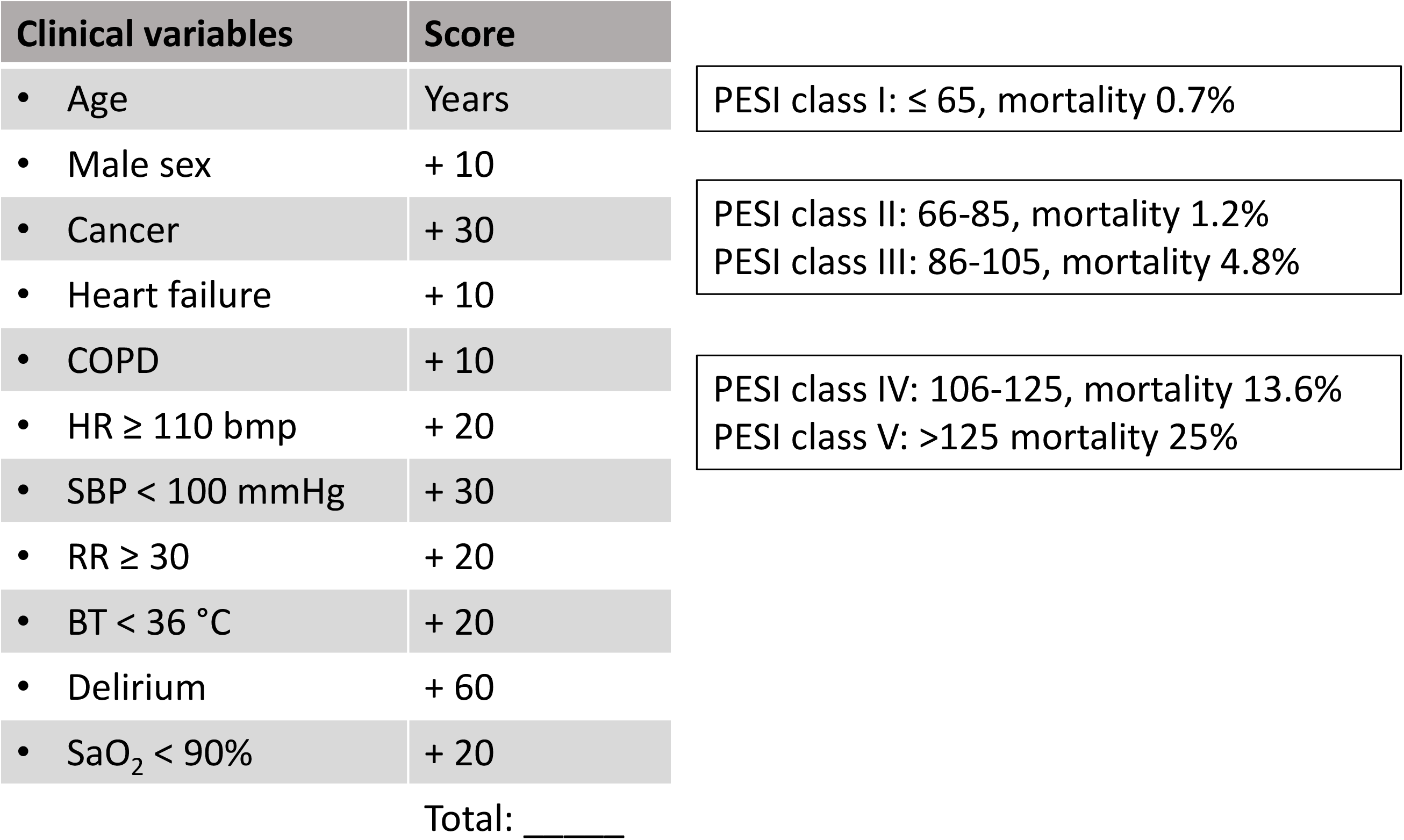
Clinical variables determining PESI Class.

**Supplemental Figure 2:**
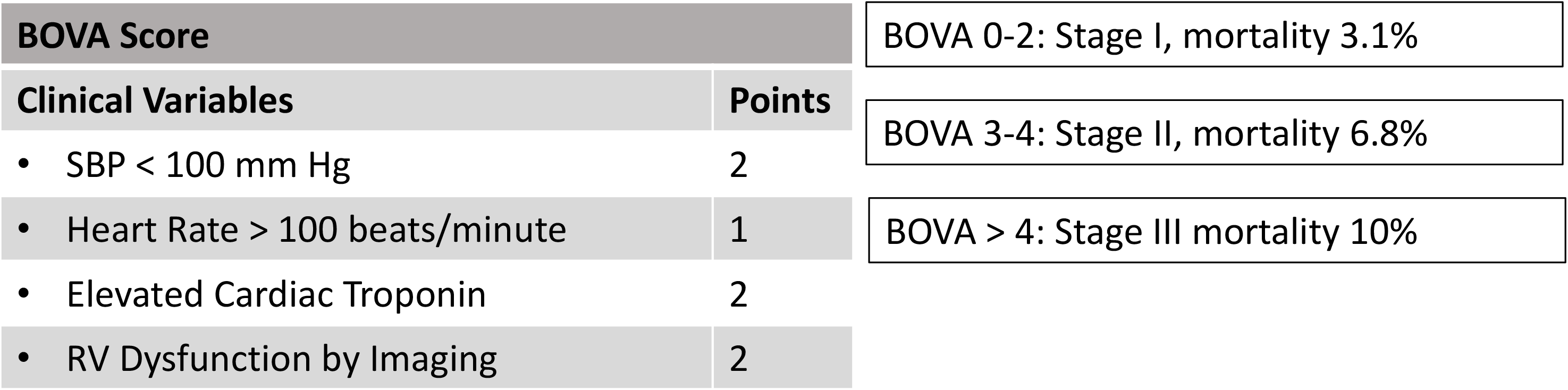
Clinical variables determining BOVA Group.

**Supplemental Figure 3:**
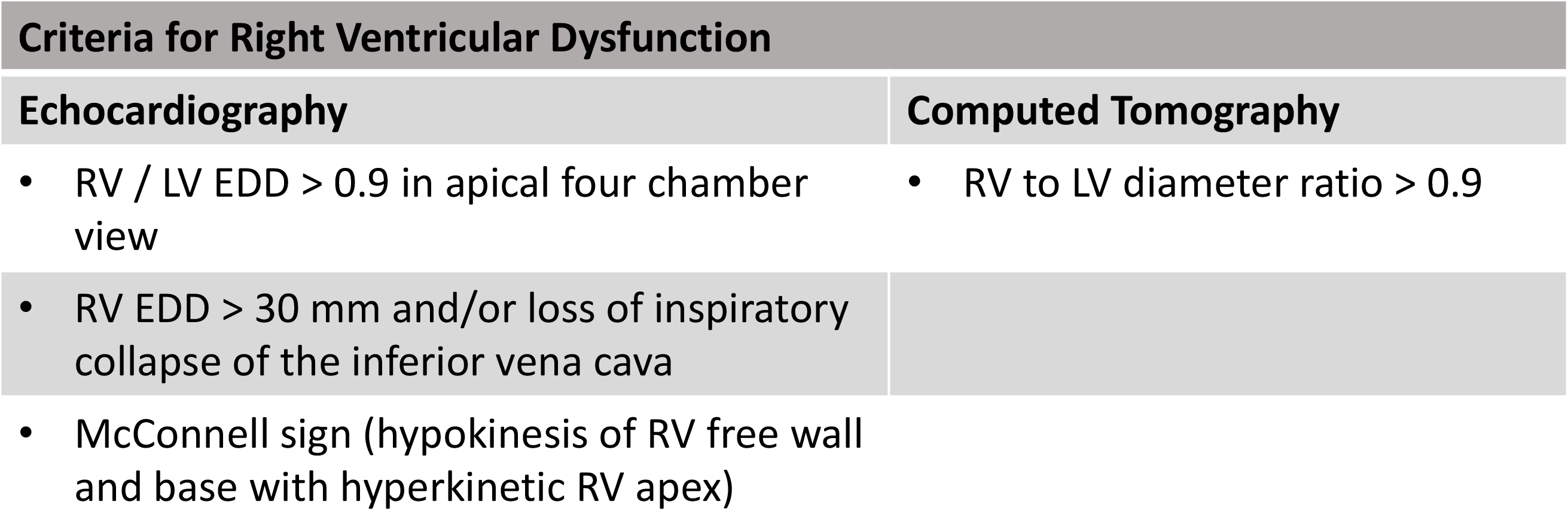
Radiographic determinants of RV dysfunction.

## References

1. Wood KE. Major pulmonary embolism: review of a pathophysiologic approach to the golden hour of hemodynamically significant pulmonary embolism. Chest. 2002; 121: 877–905.

2. Friedman T, Winokur RS, Quencer KB and Madoff DC. Patient Assessment: Clinical Presentation, Imaging Diagnosis, Risk Stratification, and the Role of Pulmonary Embolism Response Team. Semin Intervent Radiol. 2018; 35: 116–21.

3. Jimenez D, Yusen RD, Otero R, et al. Prognostic models for selecting patients with acute pulmonary embolism for initial outpatient therapy. Chest. 2007; 132: 24–30.

4. Aujesky D, Obrosky DS, Stone RA, et al. Derivation and validation of a prognostic model for pulmonary embolism. Am J Respir Crit Care Med. 2005; 172: 1041–6.

5. Aujesky D, Roy PM, Le Manach CP, et al. Validation of a model to predict adverse outcomes in patients with pulmonary embolism. Eur Heart J. 2006; 27: 476-81.

6. Aujesky D, Perrier A, Roy PM, et al. Validation of a clinical prognostic model to identify low-risk patients with pulmonary embolism. J Intern Med. 2007; 261: 597–604.

7. Masotti L, Righini M, Vuilleumier N, et al. Prognostic stratification of acute pulmonary embolism: focus on clinical aspects, imaging, and biomarkers. Vasc Health Risk Manag. 2009; 5: 567–75.

8. Bova C, Sanchez O, Prandoni P, et al. Identification of intermediate-risk patients with acute symptomatic pulmonary embolism. Eur Respir J. 2014; 44: 694–703.

9. Bikdeli B, Lobo JL, Jimenez D, et al. Early Use of Echocardiography in Patients With Acute Pulmonary Embolism: Findings From the RIETE Registry. J Am Heart Assoc. 2018; 7: e009042.

10. Kucher N, Rossi E, De Rosa M and Goldhaber SZ. Prognostic role of echocardiography among patients with acute pulmonary embolism and a systolic arterial pressure of 90 mm Hg or higher. Arch Intern Med. 2005; 165: 1777–81.

11. Cho JH, Kutti Sridharan G, Kim SH, et al. Right ventricular dysfunction as an echocardiographic prognostic factor in hemodynamically stable patients with acute pulmonary embolism: a meta-analysis. BMC Cardiovasc Disord. 2014; 14: 64.

12. Darwish OS, Mahayni A, Patel M and Amin A. Cardiac Troponins in Low-Risk Pulmonary Embolism Patients: A Systematic Review and Meta-Analysis. J Hosp Med. 2018; 13: 706–12.

13. Kearon C, Akl EA, Ornelas J, et al. Antithrombotic Therapy for VTE Disease: CHEST Guideline and Expert Panel Report. Chest. 2016; 149: 315–52.

14. Konstantinides SV, Torbicki A, Agnelli G, et al. 2014 ESC guidelines on the diagnosis and management of acute pulmonary embolism. Eur Heart J. 2014; 35: 3033–69, 69a–69k.

15. Rosovsky R, Chang Y, Rosenfield K, et al. Changes in treatment and outcomes after creation of a pulmonary embolism response team (PERT), a 10-year analysis. J Thromb Thrombolysis. 2019; 47: 31–40.

16. Xenos ES, Davis GA, He Q, Green A and Smyth SS. The implementation of a pulmonary embolism response team in the management of intermediate- or high-risk pulmonary embolism. J Vasc Surg Venous Lymphat Disord. 2019.

17. Mahar JH, Haddadin I, Sadana D, et al. A pulmonary embolism response team (PERT) approach: initial experience from the Cleveland Clinic. J Thromb Thrombolysis. 2018; 46: 186–92.

18. Elbadawi A, Wright C, Patel D, et al. The impact of a multi-specialty team for high risk pulmonary embolism on resident and fellow education. Vasc Med. 2018; 23: 372–6.

19. Secemsky E, Chang Y, Jain CC, et al. Contemporary Management and Outcomes of Patients with Massive and Submassive Pulmonary Embolism. Am J Med. 2018; 131: 1506–14 e0.

20. Jaff MR, McMurtry MS, Archer SL, et al. Management of massive and submassive pulmonary embolism, iliofemoral deep vein thrombosis, and chronic thromboembolic pulmonary hypertension: a scientific statement from the American Heart Association. Circulation. 2011; 123: 1788–830.

21. Sanchez O, Trinquart L, Colombet I, et al. Prognostic value of right ventricular dysfunction in patients with haemodynamically stable pulmonary embolism: a systematic review. Eur Heart J. 2008; 29: 1569–77.

22. Chan CM, Woods CJ and Shorr AF. Comparing the pulmonary embolism severity index and the prognosis in pulmonary embolism scores as risk stratification tools. J Hosp Med. 2012; 7: 22–7.

23. Vanderheyden M, Vrints C, Verstreken S, Bartunek J, Beunk J and Goethals M. B-type natriuretic peptide as a marker of heart failure: new insights from biochemistry and clinical implications. Biomark Med. 2010; 4: 315–20.

24. Mair J. Biochemistry of B-type natriuretic peptide--where are we now? Clin Chem Lab Med. 2008; 46: 1507–14.

25. Goetze JP. Biochemistry of pro-B-type natriuretic peptide-derived peptides: the endocrine heart revisited. Clin Chem. 2004; 50: 1503–10.

26. Semenov AG and Seferian KR. Biochemistry of the human B-type natriuretic peptide precursor and molecular aspects of its processing. Clin Chim Acta. 2011; 412: 850–60.

27. Panteghini M and Clerico A. Understanding the clinical biochemistry of N-terminal pro-B-type natriuretic peptide: the prerequisite for its optimal clinical use. Clin Lab. 2004; 50: 325–31.

28. Arbustini E, Pucci A, Grasso M, et al. Expression of natriuretic peptide in ventricular myocardium of failing human hearts and its correlation with the severity of clinical and hemodynamic impairment. Am J Cardiol. 1990; 66: 973–80.

29. Klok FA, Mos IC and Huisman MV. Brain-type natriuretic peptide levels in the prediction of adverse outcome in patients with pulmonary embolism: a systematic review and meta-analysis. Am J Respir Crit Care Med. 2008; 178: 425–30.

30. Jimenez D, Uresandi F, Otero R, et al. Troponin-based risk stratification of patients with acute nonmassive pulmonary embolism: systematic review and metaanalysis. Chest. 2009; 136: 974–82.

31. Becattini C, Vedovati MC and Agnelli G. Prognostic value of troponins in acute pulmonary embolism: a meta-analysis. Circulation. 2007; 116: 427–33.

32. Ferrando-Castagnetto F, Ricca-Mallada R, Selios V and Ferrando R. Atrial Arrhythmias and Scintigraphic “D-shape” Sign in Pulmonary Artery Hypertension. World J Nucl Med. 2017; 16: 75–7.

33. Movahed MR. D-shaped left ventricle seen on gated single-photon emission computed tomography is suggestive of right ventricular overload: the so-called Movahed’s sign. Am J Med. 2014; 127: e37.

34. Lodato JA, Ward RP and Lang RM. Echocardiographic predictors of pulmonary embolism in patients referred for helical CT. Echocardiography. 2008; 25: 584–90.

